# Sensitivity of Neurons in the Middle Temporal Area of Marmoset Monkeys to Random Dot Motion

**DOI:** 10.1101/104117

**Authors:** Tristan A. Chaplin, Benjamin J. Allitt, Maureen A. Hagan, Nicholas S. Price, Ramesh Rajan, Marcello G.P. Rosa, Leo L. Lui

**Affiliations:** Neuroscience Program, Biomedicine Discovery Institute and Department of Physiology, Monash University, Clayton, VIC 3800, Australia; ARC Centre of Excellence for Integrative Brain Function, Monash University Node, VIC 3800, Australia

**Keywords:** marmoset, Middle Temporal Area, motion, random-dot, multi-electrode arrays

## Abstract

Neurons in the Middle Temporal area (MT) of the primate cerebral cortex respond to moving visual stimuli. The sensitivity of MT neurons to motion signals can be characterized by using random-dot stimuli, in which the strength of the motion signal is manipulated by adding different levels of noise (elements that move in random directions). In macaques, this has allowed the calculation of “neurometric” thresholds. We characterized the responses of MT neurons in sufentanil/nitrous oxide anesthetized marmoset monkeys, a species which has attracted considerable recent interest as an animal model for vision research. We found that MT neurons show a wide range of neurometric thresholds, and that the responses of the most sensitive neurons could account for the behavioral performance of macaques and humans. We also investigated factors that contributed to the wide range of observed thresholds. The difference in firing rate between responses to motion in the preferred and null directions was the most effective predictor of neurometric threshold, whereas the direction tuning bandwidth had no correlation with the threshold. We also showed that it is possible to obtain reliable estimates of neurometric thresholds using stimuli that were not highly optimized for each neuron, as is often necessary when recording from large populations of neurons with different receptive field concurrently, as was the case in this study. These results demonstrate that marmoset MT shows an essential physiological similarity to macaque MT, and suggest that its neurons are capable of representing motion signals that allow for comparable motion-in-noise judgments.

**New and Noteworthy:** We report the activity of neurons in marmoset MT in response to random-dot motion stimuli of varying coherence. The information carried by individual MT neurons was comparable to that of the macaque, and that the maximum firing rates were a strong predictor of sensitivity. Our study provides key information regarding the neural basis of motion perception in the marmoset, a small primate species that is becoming increasingly popular as an experimental model.

## Introduction

The study of the neurophysiological correlates of motion perception has proven to be one the most powerful methodologies for understanding the relationship between neuronal activity and perception. Motion can be easily parameterized in terms of direction and speed, and, using random dot stimuli (Newsome et al. 1989; Britten et al. 1992; Pilly and Seitz 2009), noise can be introduced in a manner that allows for meaningful comparisons between the responses of neurons and perception (Parker and Newsome 1998). Such physiological studies have targeted the middle temporal area (MT) of the cerebral cortex, where the majority of neurons are direction selective (Allman and Kaas 1971; Dubner and Zeki 1971; Maunsell and Van Essen 1983; Albright 1984). Britten et al. (1992, 1996) used the spiking information obtained from MT neurons to decode the direction of noisy moving stimuli, and found that a substantial number of MT neurons show sensitivity that is similar to those of human and monkey observers. However, when comparable time scales were considered, individual neuronal thresholds proved to be generally higher than behavioral thresholds (Law and Gold 2008; Cohen and Newsome 2009), which is compatible with the notion that the activity of many cells needs to be combined to account for perception.

All studies to date that have investigated the sensitivity of MT neurons to random dot stimuli have been performed in macaque monkeys. However, there is now a growing interest in the marmoset, a small New World monkey with characteristics that facilitate some types of experiments that are not easily achievable in macaques. Firstly, marmosets reproduce and reach maturity relatively quickly, and are amenable to genetic modification techniques (Sasaki et al. 2009; Okano et al. 2012) and studies that manipulate development (Yu et al. 2013). Secondly, the marmoset cerebral cortex contains few sulci compared to macaques, with most visual areas, including MT, fully exposed on the outer surface of the brain (Rosa and Elston, 1998; Solomon and Rosa, 2014). This allows the use of planar and laminar electrode arrays and imaging techniques to record from large populations of neurons (Sadakane et al. 2015; Solomon et al. 2015; Zavitz et al. 2016)(Sadakane et al. 2015; Solomon et al. 2015; Zavitz et al. 2016)(Sadakane et al. 2015; Solomon et al. 2015; Zavitz et al. 2016). Furthermore, it has recently been shown that marmosets can be trained to perform visual discrimination tasks while controlling eye movements (Mitchell et al. 2014) opening the door for the recording of a large number of MT neurons while monkeys are engaged in a motion discrimination task.

Many features of marmoset MT, such as heavy myelination, first-order retinotopic map and high incidence of direction selective neurons are common to all primates (Rosa and Elston 1998). However, in marmosets, area MT is 4–5 times smaller than in macaques, and some details of response properties, including receptive field sizes and preferred spatial frequencies, are different (see Lui and Rosa 2015 for review). Whether these differences affect the ability of marmoset MT neurons to segregate signal from noise in random dot motion has yet to be tested. Moreover, the reason some MT neurons have lower neurometric thresholds than others has not yet been explored in detail, in any primate species. Using a large sample of neurons, obtained using either sequential recordings with single electrodes or parallel recordings with arrays, we explored these questions.

## Methods

### Animals and surgical preparation

Single-unit and multi-unit extracellular recordings were obtained from 11 marmoset monkeys in area MT. Several of these animals were also used for unrelated anatomical tracing and auditory physiology experiments. Experiments were conducted in accordance with the Australian Code of Practice for the Care and Use of Animals for Scientific Purposes, and all procedures were approved by the Monash University Animal Ethics Experimentation Committee.

The preparation for electrophysiology studies of marmosets has been described previously (Bourne and Rosa 2003, updated as in Yu and Rosa 2010). Briefly, anesthesia was induced with alfaxalone (Alfaxan, 8 mg/kg), allowing a tracheotomy, vein cannulation and craniotomy to be performed. After all surgical procedures were completed, the animal was administered an intravenous infusion of pancuronium bromide (0.1 mg/kgh; Organon, Sydney, Australia) combined with sufentanil (6–8 μ/kg/h, adjusted to ensure no physiological responses to noxious stimuli; Janssen-Cilag, Sydney, Australia) and dexamethasone (0.4 mg/kg/h; David Bull, Melbourne, Australia), and was artificially ventilated with a gaseous mixture of nitrous oxide and oxygen (7:3). The electrocardiogram and level of cortical spontaneous activity were continuously monitored. Administration of atropine (1%) and phenylephrine hydrochloride (10%) eye drops (Sigma Pharmaceuticals, Melbourne, Australia) resulted in mydriasis and cycloplegia. Appropriate focus and protection of the corneas from desiccation were achieved by means of hard contact lenses selected by retinoscopy.

### Electrophysiology, data acquisition and pre-processing

We used three different types of electrodes – single parylene-coated tungsten microelectrodes with exposed tips of 10 μm (in 3 marmosets; Microprobe, MD), a grid-like “Utah” electrode array (in 1 marmoset, Blackrock Microsystems, UT) and single shaft linear arrays (in 7 marmosets; NeuroNexus, MI). Single electrode recordings were made at intervals of at least 100 μm. The Utah array implant, consisting of 96 electrodes arranged in a 10×10 grid with each electrode separated by 400 μm, covered approximately two thirds of MT. The linear arrays consisted of 32 electrodes separated by 50 μm. MT recording sites were identified during experiments using anatomical landmarks and the retinotopic map (Rosa and Elston 1998), and were confirmed post mortem by histological examination.

Electrophysiological data were recorded using a Cereplex system (Blackrock Microsystems, MD) with a sampling rate of 30 kHz. Each channel was high-pass filtered at 750 Hz and spikes initially identified based on threshold crossings. Units were sorted offline using Offline Sorter (Plexon Inc., TX). Units were classified as single-units if they showed good separation on the (2 component) principal component analysis plot, and were confirmed by inspection of the inter-spike interval histogram and consistency of waveform over time. Any remaining threshold crossings were classified as multi-unit activity. The minimum isolation distance (Harris et al. 2001) of single-units was 17, with a median of 31, similar to recent studies (Lin et al. 2015; Ghodrati et al. 2016). The median spontaneous firing rate (to a blank, black screen) of multi-units was 2.1 spikes/s, indicating multi-units consisted of relatively few single-units.

Neurons were considered to be responsive if the stimulus-evoked activity was significantly greater than the spontaneous rate (Wilcoxon rank sum test, p<0.05). We excluded five single-units from adjacent channels from the linear array dataset since it was apparent they were duplicated across two channels, based on their sharp cross correlogram peak and high signal correlation (Bair et al. 2001).

### Stimuli

Stimuli were presented on a VIEWPixx3D monitor (1920 × 1080 pixels; 520 × 295 mm; 120 Hz refresh rate, VPixx Technologies) positioned 0.35 to 0.45 m from the animal on an angle to accommodate the size and eccentricity of the receptive field(s), typically subtending 70° in azimuth, and 40° in elevation. All stimuli were generated with MATLAB using Psychtoolbox-3 (Brainard 1997).

Stimuli consisted of random dots presented either in circular apertures or full screen. White Dots (106 cd/m^2^) of 0.2° in diameter were displayed on a black (0.25 cd/m^2^) background (full contrast). The density was such that there were on average 0.5 dots per °^2^, and was chosen because these parameters elicited good responses from marmoset MT when displayed on LCD monitors (Solomon et al. 2011; Zavitz et al. 2016). Dot coherence was controlled using the white noise method (i.e. Britten et al., 1992, 1996; see Pilly and Seitz 2009) by randomly choosing a subset of “noise” dots on each frame, which were displaced to random positions within the stimulus aperture. The remaining “signal” dots were moved in the same direction with a fixed displacement.

### Determination of receptive fields

Receptive fields were quantitatively mapped using a grid of either static flashed squares or small apertures of briefly presented moving dots. For single electrode recordings, stimuli were restricted to the excitatory receptive field; for array recordings, stimuli covered the full screen, so as to cover as many neurons’ receptive fields as possible. To estimate receptive field size, we fit a 2-dimensional Gaussian to the mean firing rates of the stimulus positions by minimizing the squared error using the Matlab function *lsqcurvefit.* The size was taken as the four standard deviations of the Gaussian (i.e. two standard deviations on either side of the mean). Only receptive fields that were well fit by this function (as determined by visual inspection) were used for receptive field size analyses.

### Quantitative Tests

We conducted a series of tests to determine direction selectivity, speed tuning and neurometric thresholds. All stimuli were presented for 500 ms with an inter-stimulus interval of 1000 ms. Direction and speed tuning tests used at least 8 repeats of each stimulus type, while neurometric thresholds tests used 25 repeats.

#### Single electrode recordings

We tested for direction selectivity by presenting a circular aperture of random dots that moved in 1 of 8 equally spaced directions at 8°/s. Speed tuning was then tested using random dot stimuli with speeds of 2–128°/s moving in the preferred direction. The neurometric threshold was determined by using stimuli with motion coherences of 0–100% at a near-preferred speed, in both preferred (calculated by vector sum; Ringach et al., 2002) and null directions.

#### Multi-electrode recordings

To test for direction selectivity and speed tuning, we used 100% coherence dots with all combinations of 12 equally spaced directions and a range of speeds (1–128°/s). To determine neurometric thresholds, we used full-screen dots with 12 possible directions, a speed of 20°/s and motion coherences of 0–100%. Rather than optimizing direction for each unit, as was the case in the single electrode condition, the axis of motion that was closest to the preferred/null was used for both these analyses. Since the directions used were separated by 30°, the neuron’s “true” preferred direction is at most 15° from the direction used, which is small compared to the typical direction tuning bandwidth of MT neurons (Lui and Rosa 2015). Furthermore, in single electrode experiments (both previous studies and ours) that use the “exact” preferred direction, the preferred direction is typically determined by a brief direction tuning test that is likely to have a margin of error.

### Data Analysis

#### Direction Tuning

The preferred direction was calculated using a vector sum of normalized above-spontaneous spiking rates (Ringach et al. 2002). We used four approaches to quantify direction selectivity. First, we calculated a direction index (DI, Albright 1984):

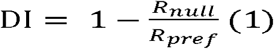

where R_pref_ and R_null_ are the above-spontaneous spike rates in the preferred and null directions respectively. DI is greater than 0 and generally less than 2, with values closer to 1 indicating stronger direction selectivity. Second, circular variance (CV) was calculated as 1 minus the length of the vector sum of normalized above-spontaneous spiking rates (Ringach et al. 2002). Third, neurons were classified as direction selective using the Rayleigh test (p<0.05). Finally, tuning bandwidth was calculated using the standard deviation of a Gaussian function fit to the direction tuning curve using least squares regression (responses of an example neuron and with fitting is shown in Figure 1A). This function included a vertical offset parameter, which effectively removes responses to motion in the null direction. As in Nover et al. (2005), the curve was fit with the complete set of trial spike rates, rather than the means, and we fitted the square root of the spikes rates to the square root the Gaussian function to homogenize the variance of spike rates (Prince et al. 2002). Only neurons that were classified as direction selective using the Rayleigh test and whose Gaussian fit R^2^ value exceeded 0.8 (as used in previous studies e.g. Nover et al. 2005, also confirmed by visual inspection of each individual fit) were included for bandwidth analyses.

**Figure 1:**
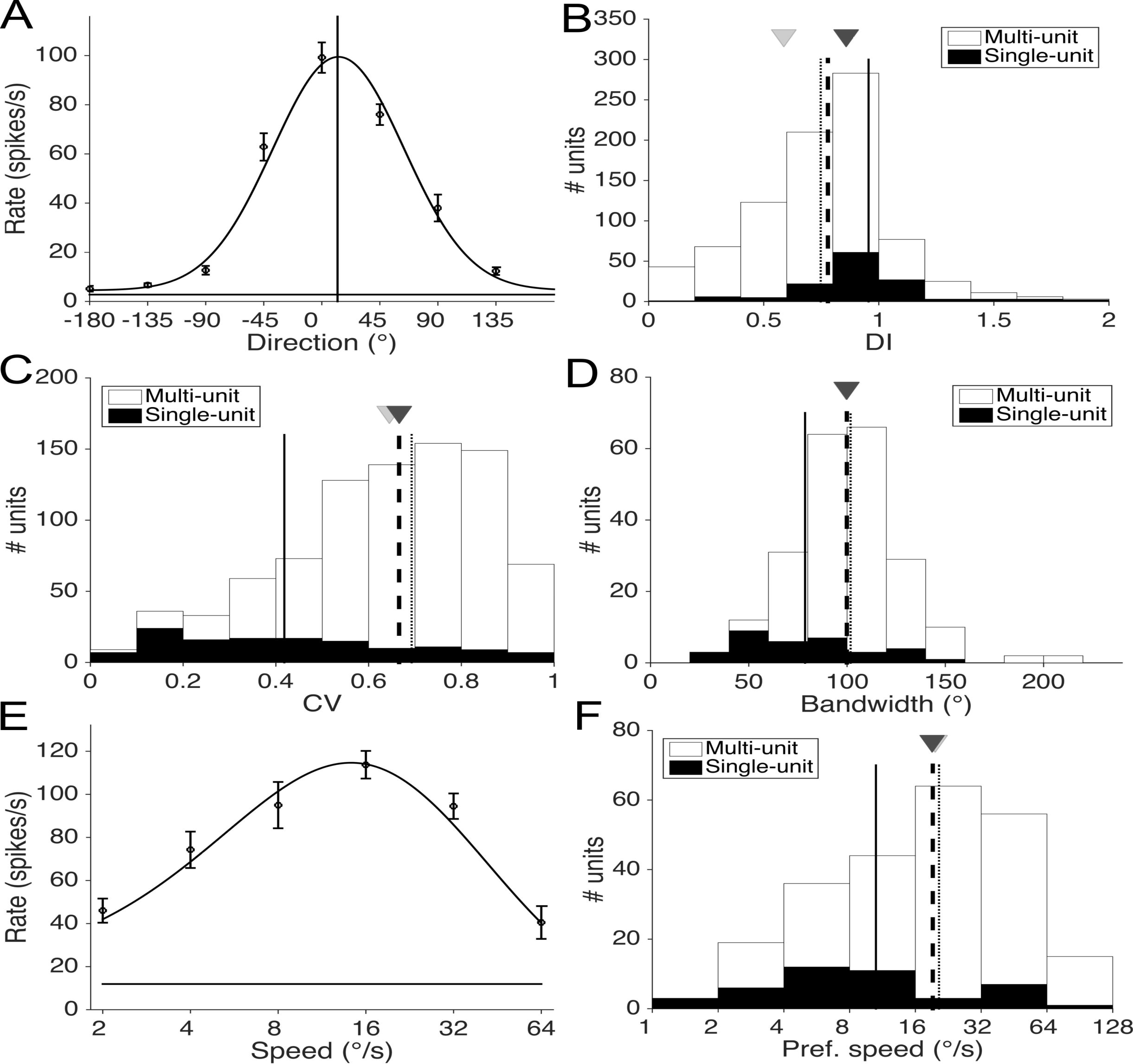
Responses of MT neurons to moving stimuli. A: Example direction tuning plot showing response with respect to direction. The highest firing rate is centered at 0 degrees and the best fitting Gaussian function to determine the tuning bandwidth is also displayed. Vertical lines show the preferred direction, error bars are standard error of the mean. B: Stacked histogram showing the distribution of direction index (DI) for singleunits (black) and multi-units (white). The medians of the overall population, single-units and the multi-units are shown as dashed, solid and dotted lines respectively. The medians of the single electrode and array recordings are shown by the dark and light grey triangles respectively. C: Distribution of circular variance (CV) of all neurons, the medians of the population and sub-groups are shown in the same convention as A. D: Distribution of direction tuning bandwidth of direction selective neurons, medians are shown in the same convention as A. E: Example speed tuning plot. The best-fitting lognormal function (Nover et al. 2005) is shown, these were used to determine the preferred speed of the neuron (the peak of the function). Error bars are standard error of the mean. F: Distribution of preferred speeds of speed tuned neurons, medians are shown in the same convention as A.

#### Speed Tuning

Speed tuning data was fit with a lognormal function (Nover et al. 2005) of the form:

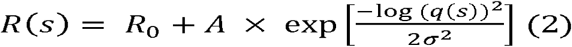

where

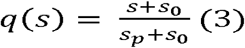

R_0_, A, s_0_, s_p_ and σ were free parameters. The preferred speed was given by s_p_, σ was the width of the speed tuning and s_0_ was an offset to fit the zero speed condition. The R0 parameter provided a vertical offset and A is a scaling parameter.

Curves were fit by minimizing the squared error using the Matlab function *lsqcurvefit.* Similar to the direction tuning fits, the speed tuning curve was fitted with the complete set of trial spike rates, rather than the means, and we fitted the square root of the spike rates to the square root of Equation 2. Only neurons whose lognormal fit R^2^ value exceeded 0.8 (as used in previous studies e.g. Nover et al. 2005, also confirmed by visual inspection of each individual fit) and whose preferred speed was within the range of speeds tested were included for speed analyses (Figure 1E).

#### Neurometric Thresholds

In order to quantify the neuron’s susceptibility to noise, we employed ideal observer analysis to determine performance of MT neurons in a direction discrimination task (Britten et al. 1992). For each level of coherence, we calculated the area under the Receiver Operator Characteristic (aROC) curve from the distributions of responses to the preferred and null directions. The aROC values were fitted using least squares regression with two variants of the Weibull function, resulting in a neurometric curve that described the neuron’s performance with respect to coherence (an aROC plot of an example neuron is shown in Figure 2B).

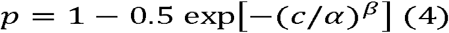

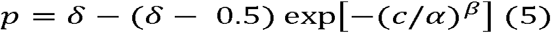

where *p* was the probability of correctly discriminating the direction of motion at coherence *c, α* was the coherence of threshold performance (p=0.82, convention established by Britten et al., 1992), β controls the slope and δ was the asymptotic level of performance (less than 1). Since Equation 5 has an extra free parameter, we used an F-test to decide whether to reject the use of the Equation 5 over Equation 4. The α was limited to between 0 and 3, the β was limited to be between 0 and 10, and the δ was limited be between 0 and 1. Neurons that did not have an aROC of at least 0.82 at 100% coherence cannot have a threshold (i.e. p(c=100)<0.82), and were excluded from analyses of thresholds, as was any neuron whose threshold exceeded 100% (since curving fitting does not guarantee the function will fit all data points perfectly).

**Figure 2:**
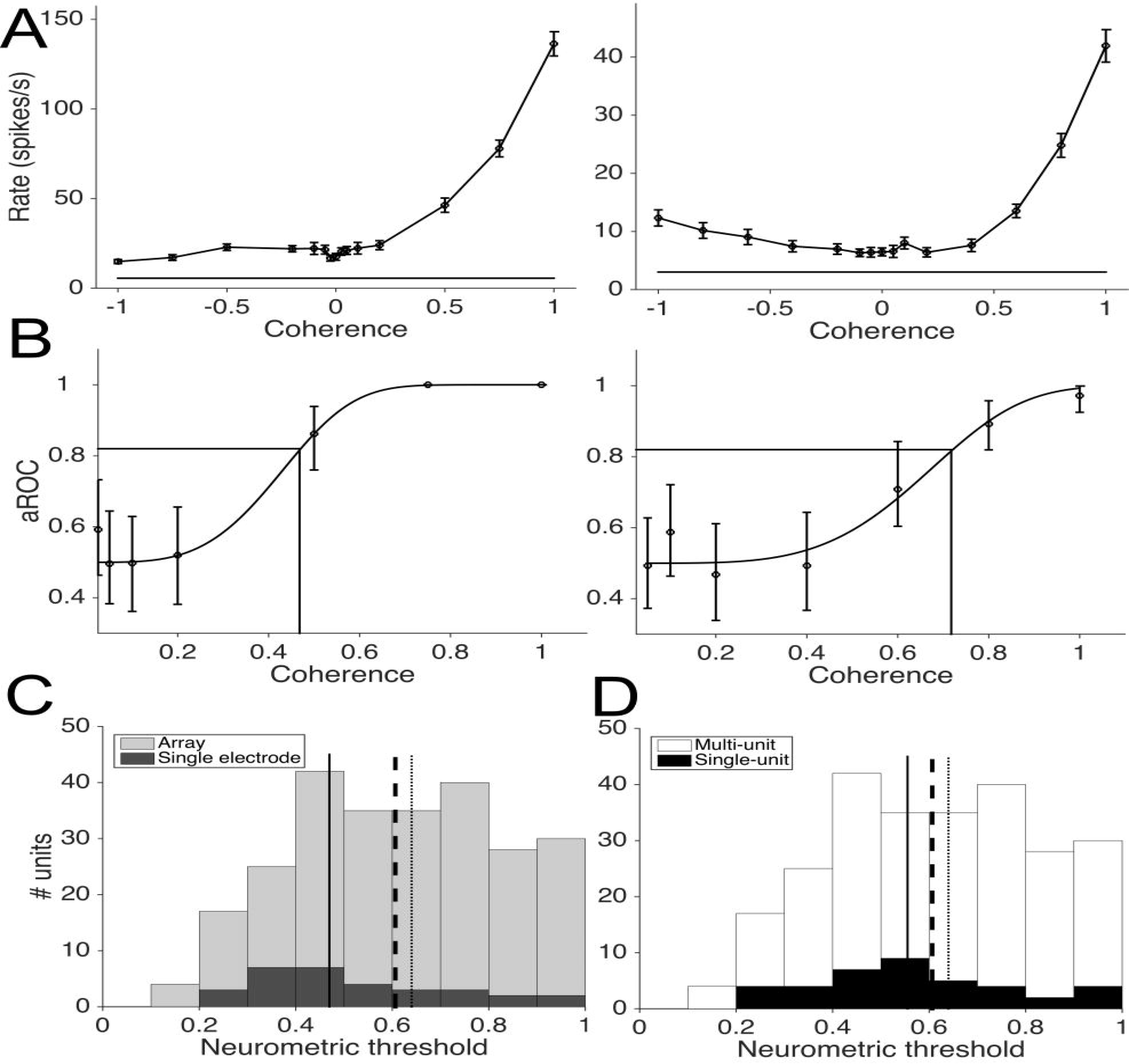
A: Responses of one neuron (left panel) and a second example neuron (right panel) to varying levels of coherence in the preferred (positive x-axis) and null directions (negative x-axis), error bars are standard error of the mean. B: Neurometric curves. Same neurons as A, data points represent aROC values with at each coherence level. The best fitting Weibull curves are shown, and the coherence level in which they reach an aROC of 0.82 is designated the neurometric threshold, shown as horizontal and vertical lines. Error bars are the 95% CI determined by bootstrapping. C: Stacked histogram showing the distribution of neurometric thresholds for single electrode (dark grey) and array (light grey) recordings. The medians of the overall population, single electrode recordings and array recordings are shown as dashed, solid and dotted lines respectively. There was a significant difference between the medians obtained with single electrodes and arrays (p=0.02). D: Stacked histogram showing the distribution of neurometric thresholds for single-unit (black) and multi-unit (white) recordings. The medians of the overall population, single-units and multi-units are shown as dashed, solid and dotted lines respectively. There was no significant difference between the median of the single-units and multi-units (p=0.18).

To obtain error bars for aROCs plots, we employed a bootstrapping method (Uka and DeAngelis 2003). Briefly, we resampled with replacement of the trials at each coherence level to obtain new sets of spike rates. These new set of rates were used to calculate aROCs as described previously. This procedure was repeated 1000 times and the 95% percentile of the bootstrapped aROCs was used as the error bars in plots.

#### Detection Thresholds

In order to determine how well neurons could detect motion in the preferred direction versus random motion, we calculated aROC comparing the distribution of spikes evoked by each coherence with the distribution of spikes evoked by zero coherence. As for neurometric thresholds, we fit a Weibull function to these data to determine the detection threshold.

#### Rate thresholds

Rate thresholds were determined by fitting a power function in the form of:

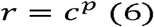

where r is the normalized firing rate of the neuron, c is the coherence, and p is a free parameter that determines the shape of the function. The p parameter was limited to be greater than zero, and thus the value of r is always zero when coherence is zero, and always equal to one when coherence is 100%. As in the direction and speed tuning curve fitting, we fit the square root of this function to the square root of each trial rate. We calculated the rate threshold by inverting the function to determine the coherence level in which the firing rate is 50% of the maximum.

#### Null aROC

The null aROC was calculated as the aROC between the zero coherence condition and the null direction 100% coherence condition. Thus a neuron whose null direction firing rate was less than the zero coherence (e.g. Figure 2A left) would have an aROC less than 0.5, whereas a neuron whose null direction firing rate was greater than the zero coherence condition (e.g. Figure 2A right) would have an aROC greater than 0.5.

#### Statistics

We used non-parametric statistics in nearly all correlations and statistical tests. Correlations were Spearman’s rho (p). Tests between two groups were made with Wilcoxon’s Rank Sum. Tests between 3 groups were made with the Kruskal-Wallis test. The exceptions to the use of non-parametric statistics were when we used a 2-way ANOVA to test for main and interaction effects of unit isolation (single-unit or multi-unit) and electrode type. When we determined the relationship between thresholds and the preferred minus null firing rates, in which we calculated the least squares line of best fit, we used Pearson’ linear correlation.

To calculate the running median of the off axis thresholds, we used sliding windows of 15 degrees in size and conducted a series of Wilcoxon’s Rank Sum tests. We started at 90 from the preferred-null axis and the direction offset was determined by offset by the first point in which the neurometric thresholds were not significantly different to the preferred direction’s neurometric thresholds.

### Histology

At the end of the recordings, the animals were given an intravenous lethal dose of sodium pentobarbitone and, following cardiac arrest, were perfused with 0.9% saline, followed by 4% paraformaldehyde in 0.1 M phosphate buffer pH 7.4. The brain was post-fixed for approximately 24 hours in the same solution, and then cryoprotected with fixative solutions containing 10%, 20%, and 30% sucrose. The brains were then frozen and sectioned into 40 μm coronal slices. Alternate series were stained for Nissl substance and myelin (Gallyas 1979). The location of recording sites was reconstructed by identifying electrode tracks, depth readings recorded during the experiment, and, in the case of single electrode penetrations, by electrolytic lesions performed at the end of penetrations. In the experiment where the Utah array was used, flat mounted sections (obtained as described in Rosa et al., 1991) were used to identify electrode locations. Finally, in experiments using linear arrays, the array shank was coated in DiI, allowing visualization under fluorescence microscope prior to staining of the sections. In coronal sections, MT is clearly identifiably by heavy myelination in the granular and infragranular layers, and in the flat mount, by its heavy myelinated oval shape (Rosa and Elston 1998). The majority of neurons reported here were histologically confirmed to be in MT, but for some penetrations in which the histology was unclear (19% of units), neurons were included on the basis of their receptive field progression and direction tuning.

## Results

### Sample size

A total of 807 recordings from MT single-units and multi-units (single electrodes: n=50; linear arrays: n=672; Utah array: n=85) were obtained. Of these, 514 (single electrodes: n=42; linear arrays: n=387; Utah array: n=85) were tested for analyses of neurometric thresholds, and 584 were used for speed selectivity tests (single electrodes: n=46; linear arrays: n=538; Utah array: n=0). For 300 units (42; 258; 0) we collected data for direction, threshold and speed.

### Marmoset MT neurons are highly direction selective

We characterized the responses of marmoset MT neurons to moving random dot stimuli in a set of equally spaced directions of motion (e.g. Figure 1A). First, we examined the direction selectivity of all responsive neurons with two measures: direction index (DI) and circular variance (CV), as these measures can be applied to any neuron (i.e. are not limited to direction selective neurons and do not require good curve fitting). DI only compares the response of the preferred and null directions responses, whereas CV gives a more general measure of direction selectivity using all directions of motion. The median DI was 0.78 (Figure 1B), and the median CV was 0.66 (Figure 1C). Next, we measured direction tuning bandwidth, which is similar to CV in that it measures the broadness of direction tuning, but unlike CV, is not affected by motion in the null direction. We measured bandwidth in a subset of neurons that were both direction selective and had good bandwidth fits (n=220), and found the median bandwidth in this subpopulation was 100° (Figure 1D). Finally, we calculated the preferred stimulus speed of speed tuned neurons (n=237, Figure 1E), and found that the distribution of preferred speeds followed a lognormal distribution, with a median of 19°/s (Figure 1F). We investigated whether single-unit or multi-units, and single electrode or array recordings impacted on any of these measures. For DI, CV and bandwidth, there was a significant main effect for isolation (i.e. single-unit versus multi-unit, 2-way ANOVA, DI: p=0.007; CV: p<0.001; bandwidth: 0.015), but no main effect was found for electrode type (DI: p=0.136; CV: p=0.061; bandwidth: p=0.303). No interaction effect was found for DI (p=0.624) and bandwidth (p=0.425), however there was a significant interaction effect for CV (p=0.04). In this case, single-units had lower CVs than multi-units in both single electrode (0.58 vs 0.66) and array recordings (0.43 vs 0.67), but the magnitude of the difference appeared to be greater in the multi-array recordings, which accounted for the interaction effect.

### Marmoset MT neurons show a wide range of neurometric thresholds

We wanted to determine marmoset MT neurons’ ability to indicate the direction of motion, i.e. how accurately an ideal observer can determine the direction of motion, from one of two opposing directions, given the spike rate from either a preferred and null direction trial. With the assumption of a hypothetical “anti-neuron”, where responses are identical in all respects, but with the preferred and null directions reversed, the aROC calculation will provide the neuron’s reliability for an ideal observer to determine opposing directions of motion (see Britten et al., 1992). When the coherence of motion was reduced, the “performance” of the neuron was invariably degraded. Figure 2A shows the responses of two example neurons to changes in coherence along the preferred axis of motion, and Figure 2B shows the corresponding aROC plots of these two neurons. We then calculated the neurometric threshold, which indicates the coherence level where the neuron’s performance was at 82%. Therefore neurons must have an aROC of at least 0.82 at 100% coherence in order to have a neurometric threshold. In our data set of 514 neurons, 260 (51%) met this criterion, of which 40 (15%) were classified as single-units (single electrodes: n=4; linear arrays: n=20; Utah array: n=16). Thresholds varied from 15% to 100% coherence, with a median of 61% coherence (Figure 2C). The median threshold obtained with single electrode recordings, in which the stimulus size and speed were optimized for each recording, was significantly lower than the median threshold from that obtained with arrays, (p=0.02 Wilcoxon Rank Sum test, medians 47% & 64% respectively, Figure 2C, dark grey and light grey bars). Although the distribution of thresholds obtained with arrays spanned a wider range (i.e. the lowest neurometric threshold recorded with an array was actually lower than that recorded with a single electrode). These differences most likely do not reflect the differences in electrode type, but the differences in experimental conditions, which includes full screen displays for the multi-electrode recordings (see Discussion). There was no statistically significant difference in thresholds between single-units and multi-units (Figure 2D, black and white bars, p=0.16), nor was there any significant interaction effect between electrode type and isolation (p=0.327, 2-way ANOVA).

### Neurometric thresholds are strongly affected by maximum firing rates

Since MT neurons displayed a wide range of neurometric thresholds, we investigated the neurophysiological factors that might predict the neurometric thresholds of MT neurons. Note that in these analyses, we are aiming to predict the neurometric threshold, which is the coherence level which the neuron achieves an aROC of 0.82, from the unit’s responses at 100% coherence.

First, we examined the relationship between firing rate and neurometric threshold. The simple mean firing rate in the preferred direction proved to be a reasonable predictor of neurometric threshold (ρ=-0.56, Figure 3A), even without taking into account the spontaneous or the null direction firing rates. However, since some neurons with high firing rates in the preferred direction also have high firing rates in the null direction, we also tested if the difference in firing rate between the preferred and null direction proved to be a stronger predictor of threshold. Moreover, to account for Poisson-like variability of neuronal responses, in which trial to trial variability scales with mean firing rate (Churchland et al. 2010), we used the difference between the square root of the preferred and null rates. This proved to be a very good predictor of neurometric threshold (ρ=0.80, Figure 3B) and accounted for over 60% of the variance in the neurometric thresholds. Thus, the neurometric threshold was predicted reliably even if only the firing rates of the preferred and null directions are known, even without testing at coherences lower than 100%. To determine if this relationship differed between electrode type and isolation, we fit a linear model to the log of the differences in firing rates of the complete dataset and to subgroups - single-units, multi-units, single electrode recordings, array recordings. We found that confidence intervals of the parameters overlapped substantially with one another, indicating that this relationship holds across these grouping (Figure 3C-D). We also compared thresholds to the ratio of the null to the preferred rate (i.e. instead of the difference) and found that it was an inferior predictor (ρ=0.60, Figure 3E).

**Figure 3:**
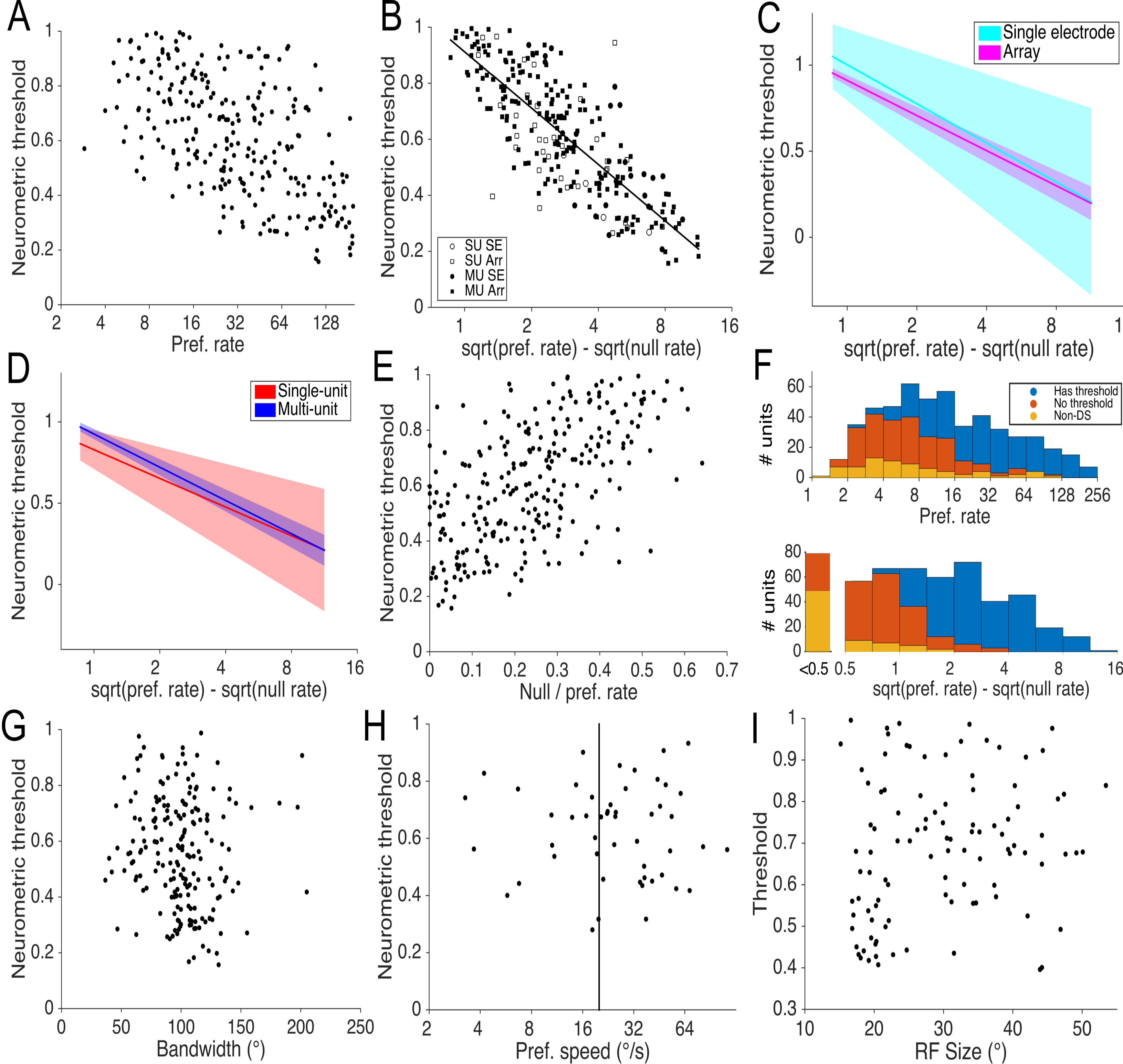
Factors affecting neurometric thresholds. A: Neurometric thresholds are shown with respect to the maximum firing rate of the neuron. Thresholds are strongly affected simply by the mean firing rate (ρ=-0.56, p<0.001). B: Neurometric thresholds are shown with respect to the difference in the square root of the firing rate of the preferred and null directions, showing a stronger relationship than in A. This relationship was well described by a logarithmic function: threshold = -0.29 log(sqrt(pref. rate) − sqrt(null rate)) + 0.9 (Pearson’s correlation r=0.81) as indicated by the line of best fit. Symbols represent whether the recording was a singleunit (SU, open) or multi-unit (MU, filled), and whether it was recorded with the single electrode (SE, circles) or array (Arr, squares). C: Linear models fit to either the single electrode (cyan) or array recordings (magenta) showing the 95% confidence interval of the fit as shading. D: Linear models fit to either the single-unit (red) or multi-unit (blue) showing the 95% confidence interval of the fit as shading. E: Neurometric threshold are shown with respect to the ratio between the null and preferred direction firing rates, showing a weaker relationship than B (ρ=-0.60, p<0.001). F: Distributions of above-spontaneous firing rates (top) and differences in the square root of the firing rate of the preferred and null directions for three classes of neuron - those with definable thresholds (blue), direction selective cells without definable thresholds (red) and non-direction selective cells (yellow, “non-DS”). Direction selective cells that did not have thresholds had both lower above spontaneous firing rates and lower differences in firing rates (p < 0.001 & p<0.001 respectively). G: Neurometric thresholds are shown with respect to direction tuning bandwidth. A small but significant relationship was found here (ρ=0.14, p= 0.047), but it was not present when controlling for difference in firing rate (ρ=-0.05, p= 0.455). H: Neurometric threshold (measured at 20°/s as indicated by black line), are shown with respect to the preferred speed of each neuron. There was no significant relationship between preferred speed and neurometric threshold (ρ=0.06, p=0.691). I: Neurometric thresholds plotted against the multi-unit recording receptive field sizes. There was a weak but significant relationship (ρ=0.22, p=0.023), which was not present when controlling for differences in firing rate (ρ=0.17, p=0.090).

The above analysis only included cells that had neurometric thresholds. Fifteen percent of cells (77/514) were excluded because they were not significantly direction selective. This proportion is similar to previous reports (see Solomon and Rosa, 2014). The remaining 177 units (34% of the data set) were direction selective neurons that were not sensitive enough for the threshold to be quantified. We extended our analyses to include these neurons, and found that both non-direction selective cells (p<0.001) and direction selective cells without thresholds (p<0.001) had lower maximal firing rates than cells with thresholds (Figure 3F). The maximum firing rates of these two groups were not significantly different from each other (p=0.543). We also investigated the difference between the square roots of the preferred and null rates. Naturally, the non-direction selective cells had significantly lower differences preferred vs null direction firing rates than the other two groups; moreover, there was a significant separation for this metric for the cells with and without thresholds (p<0.001, Figure 3F lower panel). Therefore, the difference between responses to the preferred and null directions was also a reliable predictor of whether a cell was sensitive enough to have a threshold.

We also investigated whether neurons that have sharper direction tuning (i.e. smaller tuning bandwidths) also have lower thresholds. We found only a very weak correlation between bandwidth and threshold (ρ=0.14, p= 0.047, Figure 3G), and this small effect disappeared when controlling for firing rate with a partial correlation (ρ=-0.05, p= 0.455), since there was a very weak correlation between firing rate and bandwidth (ρ=0.14, p= 0.048), i.e. neurons that have higher firing rates also tended to have broader tuning bandwidths.

Since the array threshold tests were performed using a single speed, we investigated whether the poorer neurometric thresholds came from units that preferred much slower or faster speeds than the test speed used (20°/s). For this analysis we used a subset of 50 neurons from the linear array recordings, which had both neurometric thresholds and speed tuning, and had similar preferred directions for the two sets of the tests (to ensure that the electrode position relative to the neurons had not changed over time). Figure 3H shows that neurons that preferred slow or fast speeds did not tend to have poorer thresholds than those that preferred speeds closer to 20°/s, and there was no correlation between threshold and the absolute log ratio of the preferred speed to 20°/s. (ρ=0.06, p=0.691). Therefore it is not likely that using a single speed, which in many cases was non-preferred, affected the neurometric thresholds, as long as the stimulus could drive the neurons’ responses sufficiently.

Finally, we aimed to investigate if more eccentric receptive fields had higher thresholds than those positioned closer to the fovea, given the previously described relationship between preferred speed and eccentricity (Rosa and Elston 1998). Since we did not have a precise estimate of receptive eccentricity relative to the fovea, we used receptive field size as a proxy for eccentricity since it is proportional to eccentricity (Rosa and Elston 1998). We used the receptive field size of the unsorted, multi-unit activity for which we obtained good 2D Gaussian fits (see Methods). We found only a weak correlation between receptive field size and threshold (Figure 3I, ρ=0.22, p= 0.023), and this small effect was not significant when controlling for differences in firing rate (e.g. Figure 3B) with a partial correlation (ρ=0.15, p=0.09).

In summary, we found that the difference in spike rates in response to the preferred and null directions was the most reliable predictor of the neurometric threshold, in comparison with the rate in preferred direction alone, the null-preferred spike rate ratio, and the tuning bandwidth. We also found that neurons’ sensitivity to signal within noise was relatively robust to speed relative to the neurons’ preferred speed.

### Detection thresholds are also strongly affected by maximum firing rates

Next we investigated the performance of MT neurons for detecting the presence of motion in noise. Here we asked at what coherence could MT neurons reliably detect the presence of motion in the preferred direction from the random (0% coherence) stimulus, by calculating detection thresholds (the coherence level at which the neuron’s aROC for motion versus random motion was 0.82). Among the 238 units used in this analysis, the median detection threshold was at 58% coherence. We found no difference in the medians of detection thresholds for either single-units vs multi-units or single electrode vs array recordings (p=0.79 & p=0.93 respectively), so we pooled all these recordings into one dataset. Again, the difference in firing rates was a strong predictor (ρ=-0.56, Figure 4A), while tuning bandwidth showed no correlation with detection threshold (ρ=-0.08, p=0.257).

**Figure 4:**
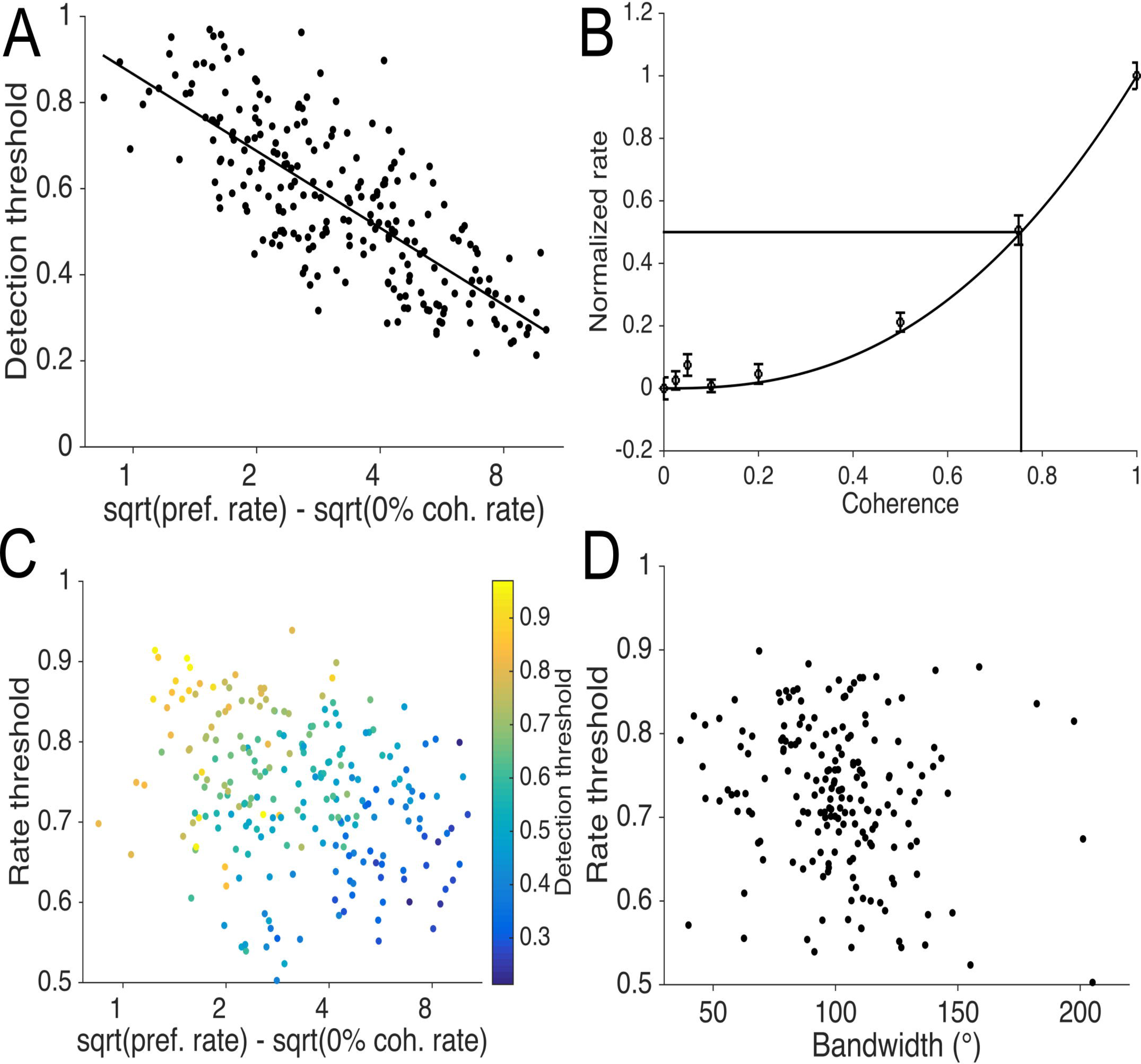
Factors affecting detection thresholds. A: Detection thresholds are shown with respect to the difference in the square root of the firing rate in the preferred direction and at zero coherence. As in Figure 5B, this relationship was well described by a logarithmic function: threshold = -0.26 log(sqrt(pref. rate) − sqrt(null rate)) + 0.86 (Pearson’s correlation r=0.77), as shown by the line of best fit. B: An example neuron demonstrating how rate thresholds were calculated. The normalized firing rate is plotted against coherence for motion in the preferred direction, error bars indicate standard error of the mean. The best fitting power function is also displayed. The rate threshold is the coherence level when the best fitting function reaches 50% of its maximum firing rate, as indicate by the black lines. C: Scatter plot showing the detection threshold (colored bar) as a combination of the rate threshold and the difference in firing rate. Neurons with the best detection thresholds have greater firing rate differences (r=-0.56, p<0.001 and lower rate thresholds (r=0.56, p<0.001). D: Rate thresholds are shown with respect to bandwidths, showing a very weak but statistically significant relationship (ρ=-0.22, p=0.002).

Since firing rates increase monotonically with increases in coherence in the preferred direction of motion (Figure 2A), we wanted to characterize the steepness of this curve in order to assess a neuron’s sensitivity to signal within noise (how the firing rate decreases with decreases in coherence) in a rate-independent manner.

In this analysis we employed a rate threshold, defined as the coherence level at which a neuron reaches 50% of the difference between the preferred direction and zero coherence firing rate (Figure 4B). Rate thresholds varied from 50% to 94% coherence with a median of 73%; hence all neurons were supra-linear (e.g. Figure 4B). Rate threshold was correlated with detection threshold (ρ=0.56), as expected. However, this result could also have been influenced by firing rates, as neurons with low rate thresholds also had larger firing rate differences (ρ=-0.25, p<0.001, Figure 4C). To test if rate threshold had any influence that was independent of firing rate, we calculated the partial correlation of the detection threshold versus the rate threshold controlled for the firing rate difference, and found that the correlation was still present (ρ=0.52, p<0.001). The combination of these two factors largely determines the detection threshold. Finally, we also tested to see if rate threshold was related to bandwidth, and found there was only a moderate correlation (ρ=-0.22, p=0.002, Figure 4D), with more broadly tuned neurons having lower rate thresholds. We also performed these analyses for subgroups of single-units vs multi-units or single electrode vs array recordings and found very similar correlations (single-unit vs multi-unit: rate difference r=-0.68 p<0.001 & r=-0.79 p<0.001, bandwidth r=-0.18 p=0.384 & r=-0.06 p=0.451; single electrode and arrays: rate difference r=-0.55 p=0.003 & r=-0.79 p<0.001, bandwidth r=0.23 p=0.305 & r=-0.11 p=0.155). Furthermore, as in Figure 3C & D, the confidence intervals for the linear fits of threshold and firing rate difference overlapped substantially.

### Responses to motion in the null direction can influence neurometric threshold

In the context of judging opposite directions of motion, we have shown above that the responses to the null direction are as important as responses to the preferred - it is the separation between the preferred and null responses that determines the neurometric threshold. We also observed that, in some neurons, the relationship between coherence and firing rate is monotonic (e.g. Figure 2A left) along the preferred axis of motion (that is, the firing rate is lower for the null direction at 100% coherence than at zero coherence), while in other neurons (e.g. Figure 2A right) there is a “U-shaped” response - the null direction produced a higher firing rate than the zero coherence. Neurons with a U-shaped coherence curve, all other factors being equal, will have higher neurometric thresholds, since the firing rate is increasing in both the preferred and null directions with increases in coherence. This would lead to a larger degree of overlap in the spike count distributions between the preferred and null directions.

We characterized this effect by calculating a “null aROC” – the aROC between the null direction and the zero coherence condition. Thus, a null aROC that is less than 0.5 is monotonic (e.g. Figure 2A left), and values greater than 0.5 represent U-shaped curves (e.g. Figure 2A right). We found a moderate correlation between null aROC and neurometric threshold (ρ=0.33, p<0.001, Figure 5A), which was still present in the partial correlation controlling for firing rate difference (ρ=0.26, p<0.001). We plotted normalized, population-averaged coherence curves (where 1 is maximal firing rate, 0 is spontaneous), and it became apparent that the U-shaped response was primarily due to multi-unit recordings from the arrays that used full screen stimuli, instead of the apertures optimized for a single RF in used single electrode recordings (Figure 5B, yellow line). We characterized the responses of aperture multi-units (n=24, 9%), full screen multi-units (n=190, 75%) and full screen single-units (n=35, 14%). The null aROCs obtained with the aperture-optimized multi-units and single-units explored with full screen stimulation were similar (Fig. 5B), but the null aROCs obtained with full screen recordings of multi-unit activity were significantly higher than both the single electrode and the singleunit array recordings (p<0.001, Kruskal-Wallis test, post hoc Rank Sum tests p=0.234, p<0.001 and p=0.031 respectively, Figure 5C). There was no significant effect of single-units/multi-units and aperture/full screen stimuli on detection of the motion in the preferred direction, with very similar detection thresholds across groups (p=0.934 Kruskal-Wallis test, Figure 5D). In contrast, it seems that the full-screen recordings suppressed the responses at and near zero coherence (Figure 5B, yellow and red lines compared to blue lines), consistent with previous findings (Hunter and Born 2011). However, it should be noted that some individual units do not follow this trend (5/35 single-units are inhibited at the zero coherence relative to the null direction and would show U-shaped coherence curves).

**Figure 5:**
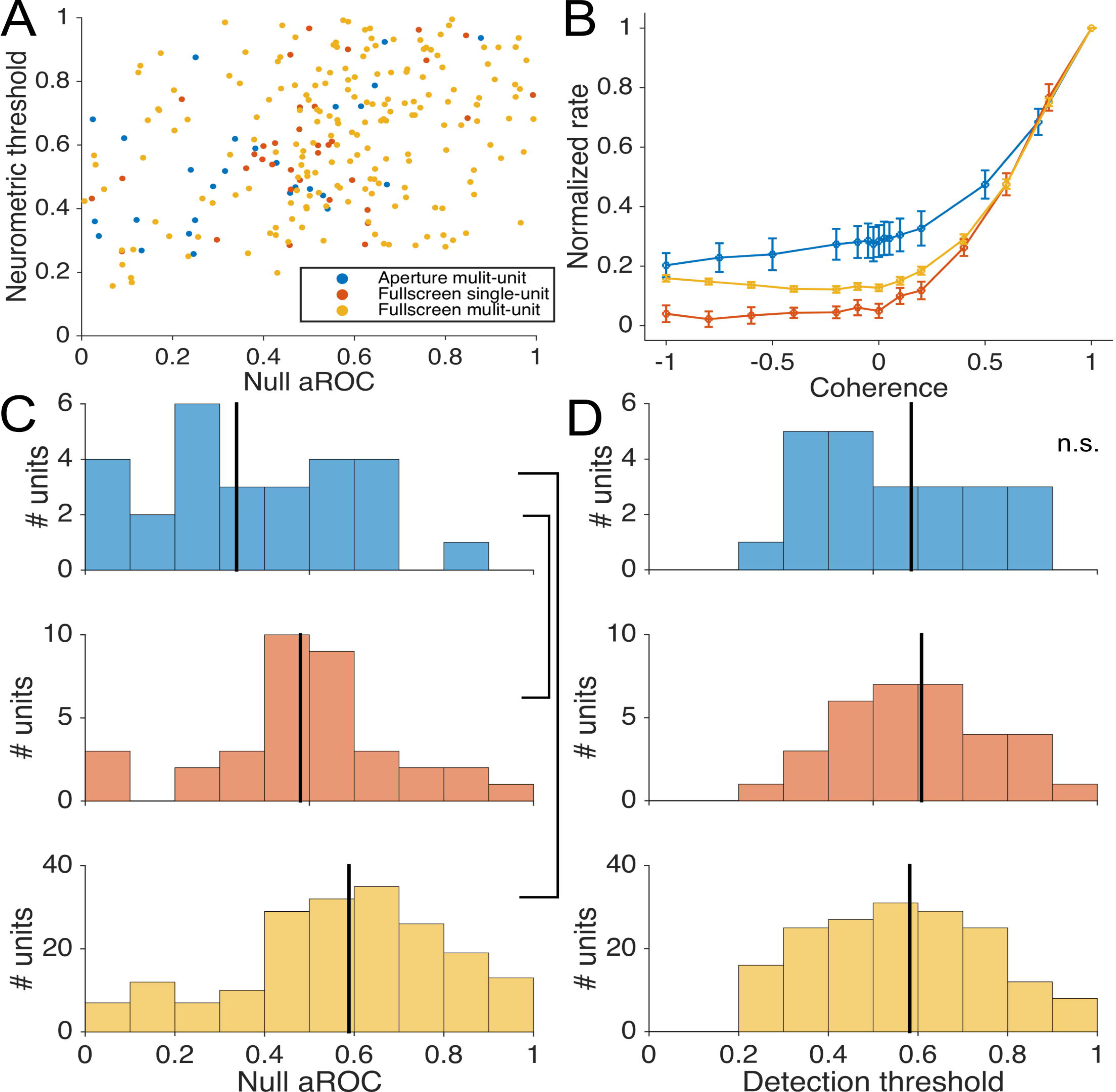
Effects of stimulus size and isolation on responses to the preferred and null direction. As indicated in legend, neurons are grouped into three types - multi-units from the single electrode in which we optimized the stimulus size (blue), single-units from the arrays in which we used full screen stimuli (red) and multi-units from the arrays in which we used full screen stimuli (yellow). A: Neurometric thresholds are shown with respect to the null aROCs for the three groupings. B: Normalized, population averaged firing rates are shown with respect to coherence for the three types of recordings. C: Distributions of the null aROCs for the three types of recordings, vertical black lines indicate medians, black lines linking histograms indicate a statistically significant difference in the medians (p<0.01). D: Distribution of detection thresholds (Figure 6A) for the three groupings, no statistically significant difference was found here (p=0.97).

### Neurometric thresholds are robust for offsets in direction of motion

The use of multi-electrode arrays allowed us to collect a large volume of data at non-optimal axes of motion. This in turn allowed us to investigate the effect of motion direction relative to the preferred-null axis on neurometric thresholds. For example, the neuron in Figure 1A has a best axis of motion centered on the 0–180° axis; however, if one were to present stimuli at the 45–225° axis, would the neuron still provide reasonably good information encoding these directions of motion? While the aROC to non-preferred directions at 100% coherence motion is predictable based on a typical MT direction tuning curve, thresholds involve changes in coherence which may affect firing rates differently at different direction offsets.

We found that the neurometric threshold generally increases for direction offsets greater than 20° (Figure 6). This was the point at which the median off-axis threshold became statistically different to the median preferred direction threshold (p<0.05). However, MT neurons were still fairly reliable at encoding direction of motion up to 60° off-axis. The maximum threshold here is limited to 300%, since it was the constraint used in the curve fitting. While this value is unrealistic (coherence cannot exceed 100%), extrapolating this parameter beyond 100% still gives an indication as to the ability of the cell to indicate opposite directions of motion. For example, a cell that is close to the 82% performance (aROC) at 100% coherence will likely to have a threshold at just over 100%, while a cells that has a much lower aROC at 100% coherence will have a threshold well beyond 100%, and perhaps towards 300% (see Britten and Newsome 1998). It can been seen in this data (as well as in Britten and Newsome 1998), that the changes in threshold with offsets in direction still follows a systematic pattern of change, despite the noisy threshold fits for some direction offsets. Furthermore, the use of a running median (instead of mean) means that the effect of not constraining the fit to 100% does not apply here until the median reaches 100% (at 60°).

We calculated the partial correlation of the off axis threshold versus the preferred axis threshold controlled for the direction offset and found a clear correlation (ρ=0.43, Figure 6, colors representing preferred axis threshold). Thus neurons with the best optimal-direction thresholds have the best off axis thresholds. We also investigated the effect of tuning bandwidth, hypothesizing that more broadly tuned neurons would have lower off axis thresholds. We found there was a weak but highly significant relationship in the partial correlation when controlling for direction offset (ρ=-0.14, p<0.001), and when controlling with both off axis direction and standard threshold (ρ=-0.16, p<0.001). These results indicate that, as expected, neurons with broad tuning bandwidths can indicate opposite directions of motion more accurately at non-optimal axes of motion than neurons with narrow tuning. However, the neurometric threshold at the unit’s preferred axis is a much better predictor of the unit’s performance at non-preferred axes.

**Figure 6:**
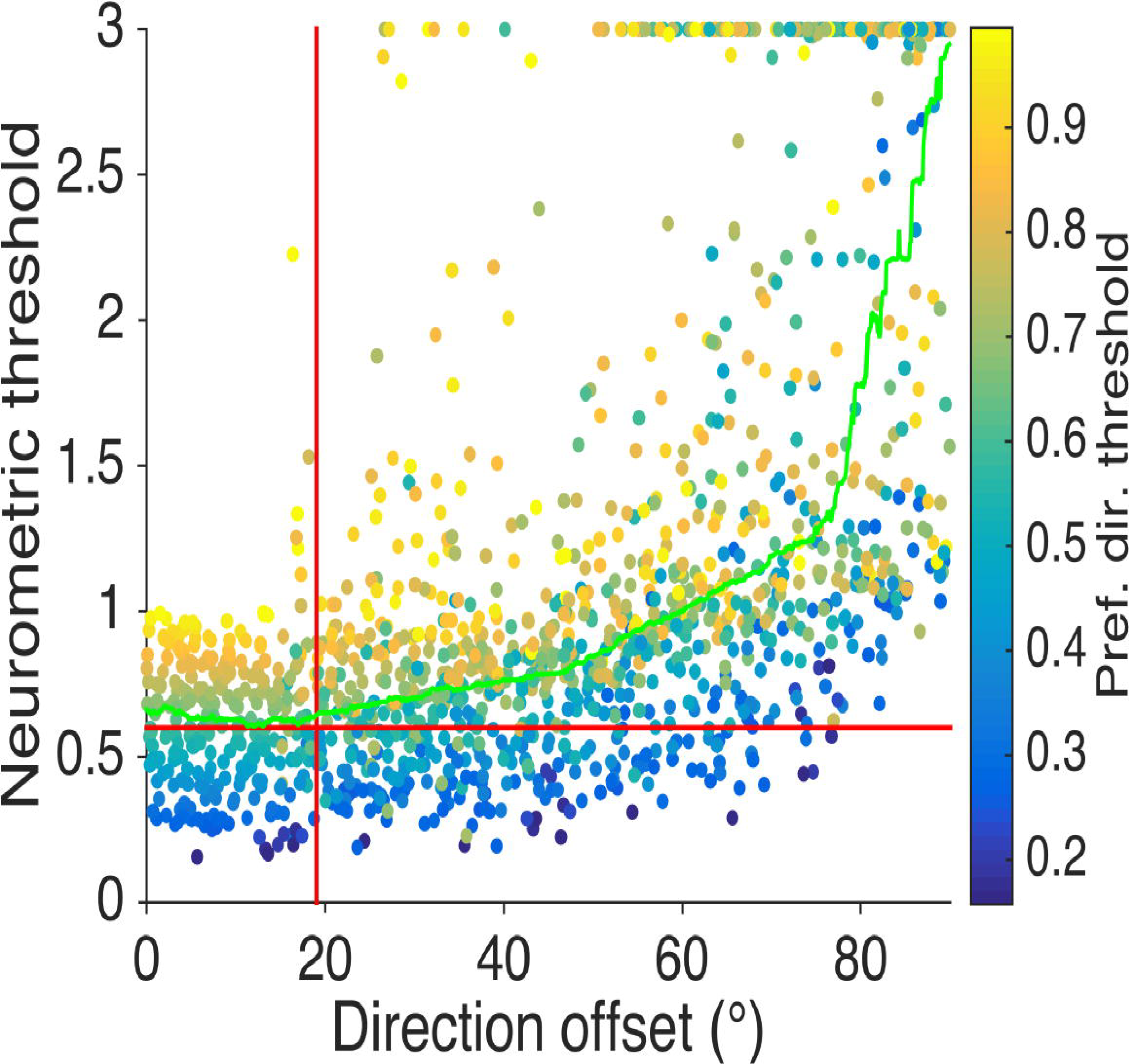
Thresholds calculated at all axes of motion, not just the preferred. Thresholds are shown with respect to the offset in direction from the preferred direction. The median optimal direction threshold is plotted as the red horizontal line, the running median is plotted in green. The direction offset in which the population median threshold is significantly different to the optimal direction threshold is plotted as a red vertical line. Data points are colored for threshold in the preferred-axis of motion.

In summary, we find here that neural sensitivity of MT neurons to signal in noise was relatively robust to changes in direction. Neurons are just as sensitive up to 20° from the preferred direction of motion, and yield useful information at least up to 60° from the preferred axis of motion.

## Discussion

We have characterized the direction selectivity of marmoset monkey MT neurons to random-dot motion stimuli. The majority of the responses were highly direction selective, as reported by previous studies in this species, using various types of stimuli (luminance and coherent motion-defined bars: Lui et al., 2012; gratings: Lui et al., 2007a; Davies et al., 2016; random dots: Solomon et al., 2011; see Lui and Rosa, 2015, for a review). Building upon these findings, we studied the extent to which the responses are influenced by different levels of random-dot motion noise. Based on an ideal observer analysis, we found that MT neurons show a wide range of sensitivities, with some neurons having a neurometric threshold as low as 15% coherence, to those neurons that could not reach unambiguous detection of the direction even in the absence of noise. Moreover, we found that the neural sensitivity could be reliably predicted simply from the responses to the preferred and null directions, while direction tuning bandwidths did not predict neurometric thresholds.

Finally, the sensitivity of many neurons was reasonably robust, allowing correct discrimination even upon presentation of non-optimal directions, speeds and stimulus sizes.

### Comparison with the macaque monkey neurometric thresholds

The macaque monkey is the only primate species for which comprehensive analyses including neurometric functions has been achieved for the sensitivity of MT neurons to motion. The increasing use of marmosets in visual neuroscience demands knowledge of the similarities, and potential differences between these species, which have evolved independently from a common ancestor for approximately 35 million years (Schrago 2007).

In the macaque, Britten et al. (1992, 1996) reported a majority of neurons with much lower neurometric thresholds than those reported here. However, those studies used much longer display times (2 seconds), which likely account for the lower neurometric thresholds. Studies that have used the “reaction time” version of the task, where monkeys essentially controlled viewing times, have yielded a variety of results. The viewing times, which varied from ≈300–750 ms, are more comparable to the data obtained in the present study. With this method, Cohen and Newsome (2009) reported a mean neurometric threshold of approximately 53%, while values obtained by other studies (Roitman and Shadlen 2002; Law and Gold 2008) were substantially lower at approximately 20%. The differences in neurometric thresholds between these studies was also reflected in their monkeys’ psychophysical performance; indeed, monkeys that perform better in the direction of motion discrimination task tend to have lower neurometric thresholds (Britten et al., 1992), and neurometric thresholds were correlated with daily fluctuations in behavioral performance. The values reported in the present study are only slightly higher than those of Cohen and Newsome (2009), thus, higher than those of Roitman and Shadlen (2002) and Law and Gold (2008), but before direct comparisons can be made, several other factors must be considered.

Multi-electrode recordings across the marmoset MT required the use of large (full screen) stimuli in order to stimulate the maximum number of receptive fields. We also choose a stimulus speed that was likely to evoke responses from the largest number of MT cells. Therefore, the size and speed of the stimulus were not always optimal for each cell. Moreover, most of our recordings were from multi-unit clusters, and were recorded using arrays. While we showed here that isolation (i.e. single-unit vs multi-unit) did not affect the neurometric thresholds, the lack of optimization for each recording site was likely to result in submaximal firing rates (and hence, affect neurometric thresholds). Our experiments were performed in anaesthetized preparations, which eliminates behavioral factors known to modulate cortical activity, such as attention (Treue and Martinez-Trujillo 1999; McAdams and Maunsell 2000). Finally, we implemented dot stimuli that were known to elicit strong responses in marmoset MT (Solomon et al. 2011; Zavitz et al. 2016), and these were not identical to those used in previous macaque studies (i.e. Britten et al., 1992). We also note here that different methods of generating noise dots have been used in previous macaque studies, the methods of implementation have implications on behavioral results (Pilly and Seitz, 2009), and thus may elicit different neural responses.

The balance of these factors suggests that the median neurometric threshold we found (63%) is an overestimate in the context of previous studies (our single electrode median was 47%). Together with the evidence of a substantial proportion of neurons with low thresholds, this would suggest, at least in principle, that marmoset MT neurons encode sufficient information about the direction of motion for the animals to perform the behavioral task. We also note here that, without doubt, perception relies on a pool of neurons, therefore, contributions from a population of weakly correlated neurons means that the animal can perform much better than most individual neurons. In macaques, psychometric thresholds were found to be 2.2 to 2.7 times less than the mean neurometric threshold of single neurons (Cohen and Newsome, 2009).

### Factors affecting neurometric and detection thresholds

We also examined a number of factors that can affect neurometric and detection thresholds. We found that the difference in the square root of firing rates (which accounts for Poisson-like variability) of the preferred and null directions at 100% coherence could quite accurately predict the ability to discriminate or detect motion in noise, accounting for more than 60% of the variance (Figure 3B). Thus, it is possible to use this metric to obtain a fairly accurate prediction of how a neuron’s aROC will change with decreases in coherence, without testing a range of coherences. Moreover, there was a large population of direction selective cells (34%) that did not have definable thresholds because the differences in the square root of preferred and null direction rates were too small (Figure 3C, lower).

Using the difference in firing rates also allowed good predictions regarding motion detection thresholds (Figure 4A). For detection thresholds, we also calculated a rate threshold in order to characterize each neuron’s response to changes in coherence in a rate-independent way (Figure 4B), but this measure is still somewhat correlated with firing rates (Figure 4C). The fact that all neurons showed supra-linear response to increases in coherence in the preferred direction makes it difficult for neurons with low (maximum) firing rates to have low neurometric thresholds, as neurons will not substantially increase their firing rates until coherences of over 50%. However, despite the fact that all neurons were supra-linear, and rate threshold spanned a narrow range, some neurons were more tolerant to noise, which impacts on the neuron’s ability to detect motion independent of spike rate. Interestingly, neurons that were more tolerant to noise also tended to have slightly broader tuning bandwidths, suggesting neurons that are better at integrating directions of motion may also be more tolerant to noise.

In macaques, neurons that carry the most task-relevant information, or the most sensitive neurons, are more correlated with behavioral choice, independent of stimulus (Britten et al., 1996). Whether these behavioral correlations may arise from neurons that contribute to the task, or through neuron-to-neuron correlations within the decision pool (see Nienborg et al., 2012), the sensitivity of each neuron presumably has to be learned downstream by the decision circuit (Law and Gold 2008). Our results suggest that a simpler strategy may be possible; the most sensitive neurons can simply be identified by their higher firing rates; thus potentially contributing more to synaptic integration downstream. Optimal read-out of population activity to form a perceptual decision posits that the neurons that carry more task relevant information have a larger contribution to the decision pool (e.g. Graf et al., 2011, Zavitz et al., 2016), and these weights could be assigned using a relatively straightforward mechanism, based on maximum firing rates. The relative homogeneity in rate threshold means that neurons that carry the most task relevant information at 100% coherence are also likely to carry the most information in situations with noise. Therefore, a coherence-invariant read-out template, with weights based on responses to the 100% coherent stimulus could, in theory, generalize to other coherences. This strategy would imply that a decision can be formed without knowing the coherence of the stimulus *a priori.*

### Comparison with the macaque monkey direction selectivity and speed tuning

At first sight, our analysis of direction selectivity seems to indicate the marmoset MT neurons are less direction selective than macaque MT. For example, measures of DI in macaque by Albright (1984) report a much higher mean value, in excess of 1, compared to our median of 0.78. However, our measurement of CV (median = 0.66) is very similar to other reports in macaques (approximately 0.7 in macaques, Cui et al., 2013). The median direction tuning bandwidth reported here was only slightly broader than what was reported in macaques by Albright (1984): 100° compared to 83°. Moreover, it should be noted that our measurements of DI and CV include all responsive units in our dataset, and the use of arrays meant that our recordings were not biased towards highly active units. Therefore our data likely constitute an unbiased description of direction selectivity in MT. Finally, while having minimal effects on the responses to the preferred direction, the full screen presentation together with multi-unit recordings may increase the responses to the null direction relative to the preferred direction (Figure 5). This could subsequently lower the DI in comparison to studies that use stimuli tailored to the size of the classical receptive field, which was the method employed by the vast majority of studies to date that examined the responses of MT cells (e.g. Albright, 1984; Britten et al., 1992, 1996; DeAngelis and Uka, 2003).

We also investigated the preferred speeds of marmoset MT neurons. In general, the range of preferred speeds was similar to those found in the macaque. However, the median optimal speed in marmosets was higher than that of reported in macaques (Nover et al. 2005), similar to results obtained with gratings (Lui et al. 2007a). The distribution of preferred speeds in the present data followed a normal distribution on a logarithmic scale (Figure 1F; see also Lui and Rosa, 2014). This is in contrast to the log-uniform distribution reported for the macaque MT, which is consistent with speed discrimination performance where just-noticeable speed differences scale with the pedestal or reference speed (Nover et al. 2005). This suggests another hypothesis to be tested behaviorally, namely that the just noticeable differences may not scale uniformly across slow, medium and fast speeds. Alternatively, these apparent differences in the distribution of speed tuning in marmosets and macaques maybe caused by differential sampling of visual space in these studies, since neurons with foveal receptive fields tend to prefer slower speeds than those set in the periphery in both species (Maunsell and Van Essen 1983; Lui et al. 2007a).

### Multi-electrode arrays vs. single contact electrodes

The use of multi-electrode recordings allowed us to collect large amounts of data on the responses at nonoptimal directions. We showed that neural sensitivity was relatively robust with respect to the direction of motion. Thresholds were not significantly higher when determining opposite directions of motion when stimuli moved in directions up to 20° off their optimal, and neurons were able to provide reliable information for directions up to 60° from optimal. This result implies that neurons with optimal directions of motion up to 60° away from the axis of motion in question can contribute to the decision pool. These results reveal slightly higher thresholds than those reported in the macaque (Britten and Newsome 1998), although, as argued above, this may reflect the exact experimental conditions. However, the rate of increase of neurometric threshold with respect to direction away from the optimal preferred-null axis is comparable in the two species. The larger data set in the present study may be more representative of the sensitivities of MT neurons to non-optimal directions.

We did not observe a significant decline in neural sensitivity with respect to the magnitude of the difference between the test speed (20°/s) and the neuron’s preferred speed, suggesting that neural sensitivity is quite robust to variations in this parameter. However, this result should be interpreted with caution. First, we chose a test speed that was approximately the mean preferred speed of MT neurons. Therefore, this result may not necessarily be generalizable to very low or high speeds. Second, neurons were included in this analysis if they were responsive at 20°/s and satisfied the inclusion criteria; thus, it remains a possibility that more neurons were excluded at due to their lack of responses at this speed. Nonetheless, our results show that MT neurons are able to contribute to direction discrimination of non-optimal direction and speed. These results have potential implications on the make-up of the neural pool that can contribute to the formation of perceptual decision.

Overall, single electrode recordings have a lower median neurometric thresholds compared to the arrays. The use of electrode arrays meant that we did not bias our recordings to large, highly active neurons (Carandini et al. 2005). This resulted in a large dataset that included many low firing rate single-units and multi-units (which may be bypassed in exploration with a single electrode due to a perceived lack of responsiveness).

Furthermore, the use of full screen stimuli to cover many receptive fields, imposed by different receptive fields during array recordings, seems to not only suppress responses to lower coherence stimuli, which is in line with a previous study (Hunter and Born 2011), but also all coherences in the null direction of single-units (Figure 5B, red line). The lower overall firing rates for the array recordings would account for the differences between the preferred and null spike rates, and in turn, the lower neurometric thresholds recorded with the single electrode. Therefore it is possible that the neurons with the lowest neurometric thresholds are disproportionally represented by those that have the least surround suppression. Also, excitation above the 0% coherence stimulus in the null direction (“U-shaped responses”) was more commonly observed for full screen multi-units (Figure 5). This could be explained by a combination of tuned and un-tuned gain normalization mechanisms, which have been demonstrated in MT (Simoncelli and Heeger 1998; Rust et al. 2006), and have been implicated in size tuning (Born and Bradley 2005), including in marmosets (Lui et al. 2007b). The balance of excitation and inhibition here depends on stimulation inside and outside the classical receptive field. Also, multi-units by definition encompass the activity of multiple neurons, the combination of which may lead to excitation in the null direction in combination with stimulation outside the classical receptive field. As the majority of our recordings with multi-electrodes were multi-units, this factor, combined with our demonstration that responses to the null direction affected the sensitivity of MT neurons, provides further explanation as to why neurons recorded with multi-electrode arrays were somewhat less sensitive to motion signals embedded in noise. Finally, even though differences in response properties were observed in responses to the stimulus conditions associated with each electrode type, the relationship between the difference in firing rate between preferred and null direction was a strong predictor of neurometric threshold regardless of these conditions.

## Acknowledgements

This project was funded by the Australian Research Council (DE130100493 to LL; CE140100007 to MR) and by the National Health and Medical Research Council of Australia (APP1066232 to LL and APP1083152 to MR). Tristan Chaplin was funded by an Australian Postgraduate Award. We thank Katrina Worthy for the histological work and Sofia Bakola for advice on manuscript. We also thank Janssen-Cilag for the donation of sufentanil citrate.

## References

Albright TD. Direction and orientation selectivity of neurons in visual area MT of the macaque. J Neurophysiol 52: 1106-1130, 1984.

Allman JM, Kaas JH. Representation of the visual field in striate and adjoining cortex of the owl monkey (Aotus trivirgatus). Brain Res 35: 89-106, 1971.

Bair W, Zohary E, Newsome WT. Correlated firing in macaque visual area MT: time scales and relationship to behavior. J Neurosci 21: 1676-97, 2001.

Born RT, Bradley DC. Structure and function of visual area MT. Annu Rev Neurosci 28: 157-189, 2005.

Bourne JA, Rosa MGP. Preparation for the in vivo recording of neuronal responses in the visual cortex of anaesthetised marmosets (Callithrix jacchus). Brain Res Protoc 11: 168-177, 2003.

Brainard DH. The Psychophysics Toolbox. Spat Vis 10: 433-6, 1997.

Britten KH, Newsome WT. Tuning bandwidths for near-threshold stimuli in area MT. J Neurophysiol 80: 762-770, 1998.

Britten KH, Newsome WT, Shadlen MN, Celebrini S, Movshon JA. A relationship between behavioral choice and the visual responses of neurons in macaque MT. Vis Neurosci 13: 87-100, 1996.

Britten KH, Shadlen MN, Newsome WT, Movshon JA. The Analysis of Visual MotionB: A Comparison of Neuronal and Psychophysical Performance. J Neurosci 12: 4745-4765, 1992.

Carandini M, Demb JB, Mante V, Tolhurst DJ, Dan Y, Olshausen Ba, Gallant JL, Rust NC. Do we know what the early visual system does? J Neurosci 25: 10577-97, 2005.

Churchland MM, Yu BM, Cunningham JP, Sugrue LP, Cohen MR, Corrado GS, Newsome WT, Clark AM, Hosseini P, Scott BB, Bradley DC, Smith MA, Kohn A, Movshon JA, Armstrong KM, Moore T, Chang SW, Snyder LH, Lisberger SG, Priebe NJ, Finn IM, Ferster D, Ryu SI, Santhanam G, Sahani M, Shenoy K V. Stimulus onset quenches neural variability: a widespread cortical phenomenon. Nat Neurosci 13: 369-378, 2010.

Cohen MR, Newsome WT. Estimates of the contribution of single neurons to perception depend on timescale and noise correlation. J Neurosci 29: 6635-48, 2009.

Cui Y, Liu LD, Khawaja FA, Pack CC, Butts DA. Diverse Suppressive Influences in Area MT and Selectivity to Complex Motion Features. J Neurosci 33: 16715-16728, 2013.

Davies AJ, Chaplin TA, Rosa MGP, Yu H-H. Natural motion trajectory enhances the coding of speed in primate extrastriate cortex. Sci Rep 6, 2016.

DeAngelis GC, Uka T. Coding of horizontal disparity and velocity by MT neurons in the alert macaque. J Neurophysiol 89: 1094-111, 2003.

Dubner R, Zeki SM. Response properties and receptive fields of cells in an anatomically defined region of the superior temporal sulcus in the monkey. Brain Res 35: 528-32, 1971.

Gallyas F. Silver staining of myelin by means of physical development. Neurol Res 1: 203-9, 1979.

Ghodrati M, Alwis DS, Price NSC. Orientation selectivity in rat primary visual cortex emerges earlier with low-contrast and high-luminance stimuli. Eur J Neurosci 44: 2759-2773, 2016.

Harris KD, Hirase H, Leinekugel X, Henze DA, Buzsáki G. Temporal interaction between single spikes and complex spike bursts in hippocampal pyramidal cells. Neuron 32: 141-149, 2001.

Hunter JN, Born RT. Stimulus-dependent modulation of suppressive influences in MT. J Neurosci 31: 678-686, 2011.

Law C-T, Gold JI. Neural correlates of perceptual learning in a sensory-motor, but not a sensory, cortical area. Nat Neurosci 11: 505-513, 2008.

Lin IC, Okun M, Carandini M, Harris KD. The Nature of Shared Cortical Variability. Neuron 87: 644–656, 2015.

Lui LL, Bourne JA, Rosa MGP. Spatial and temporal frequency selectivity of neurons in the middle temporal visual area of new world monkeys (Callithrix jacchus). Eur J Neurosci 25: 1780-92, 2007a.

Lui LL, Bourne JA, Rosa MGP. Spatial summation, end inhibition and side inhibition in the middle temporal visual area (MT). J Neurophysiol 97: 1135-48, 2007b.

Lui LL, Dobiecki AE, Bourne JA, Rosa MGP. Breaking camouflage: responses of neurons in the middle temporal area to stimuli defined by coherent motion. Eur J Neurosci 36: 2063-76, 2012.

Lui LL, Rosa MGP. Structure and function of the middle temporal visual area (MT) in the marmoset: Comparisons with the macaque monkey. Neurosci Res 93: 62-71, 2015.

Maunsell JHR, Van Essen DC. Functional properties of neurons in middle temporal visual area of the macaque monkey. I. Selectivity for stimulus direction, speed, and orientation. J Neurophysiol 49: 1127-29 47, 1983.

McAdams CJ, Maunsell JHR. Attention to Both Space and Feature Modulates Neuronal Responses in Macaque Area V4 Attention to Both Space and Feature Modulates Neuronal Responses in Macaque Area V4. J Neurophysiol 83: 1751-1755, 2000.

Mitchell JF, Reynolds JH, Miller CT. Active vision in marmosets: a model system for visual neuroscience. J Neurosci 34: 1183-94, 2014.

Newsome WT, Britten KH, Movshon JA. Neuronal correlates of a perceptual decision. Nature 341: 52–54, 1989.

Nienborg H R., Cohen M, Cumming BG. Decision-related activity in sensory neurons: correlations among neurons and with behavior. Annu Rev Neurosci 35: 463-483, 2012.

Nover H, Anderson CH, DeAngelis GC. A logarithmic, scale-invariant representation of speed in macaque middle temporal area accounts for speed discrimination performance. J Neurosci 25: 10049–60, 2005.

Okano H, Hikishima K, Iriki A, Sasaki E. The common marmoset as a novel animal model system for biomedical and neuroscience research applications. Semin Fetal Neonatal Med 17: 336-40, 2012.

Parker AJ, Newsome WT. Sense and the single neuron: probing the physiology of perception. Annu Rev Neurosci 21: 227-277, 1998.

Pilly PK, Seitz AR. What a difference a parameter makes: a psychophysical comparison of random dot motion algorithms. Vision Res 49: 1599-612, 2009.

Prince SJD, Pointon AD, Cumming BG, Parker AJ. Quantitative analysis of the responses of V1 neurons to horizontal disparity in dynamic random-dot stereograms. J Neurophysiol 87: 191-208, 2002.

Ringach DL, Shapley RM, Hawken MJ. Orientation selectivity in macaque V1: diversity and laminar dependence. J Neurosci 22: 5639-51, 2002.

Roitman JD, Shadlen MN. Response of neurons in the lateral intraparietal area during a combined visual discrimination reaction time task. J Neurosci 22: 9475-89, 2002.

Rosa MGP, Elston GN. Visuotopic organisation and neuronal response selectivity for direction of motion in visual areas of the caudal temporal lobe of the marmoset monkey (Callithrix jacchus): middle temporal area, middle temporal crescent, and surrounding cortex. J Comp Neurol 393: 505-27, 1998.

Rosa MGP, Gattass R, Soares JGM. A quantitative analysis of cytochrome oxidase-rich patches in the primary visual cortex of Cebus monkeys: topographic distribution and effects of late monocular enucleation. Exp Brain Res 84: 195-209, 1991.

Rust NC, Mante V, Simoncelli EP, Movshon JA. How MT cells analyze the motion of visual patterns. Nat Neurosci 9: 1421-1431, 2006.

Sadakane O, Masamizu Y, Watakabe A, Terada S-I, Ohtsuka M, Takaji M, Mizukami H, Ozawa K, Kawasaki H, Matsuzaki M, Yamamori T. Long-Term Two-Photon Calcium Imaging of Neuronal Populations with Subcellular Resolution in Adult Non-human Primates. Cell Rep 13: 1989-1999, 2015.

Sasaki E, Suemizu H, Shimada A, Hanazawa K, Oiwa R, Kamioka M, Tomioka I, Sotomaru Y, Hirakawa R, Eto T, Shiozawa S, Maeda T, Ito M, Ito R, Kito C, Yagihashi C, Kawai K, Miyoshi H, Tanioka Y, Tamaoki N, Habu S, Okano H, Nomura T. Generation of transgenic non-human primates with germline transmission. Nature 459: 523-7, 2009.

Schrago CG. On the time scale of new world primate diversification. Am J Phys Anthropol 132: 344-354, 2007.

Simoncelli EP, Heeger DJ. A Model of Neuronal Responses in Visual Area MT. Vision Res 38: 743-761, 1998.

Solomon SG, Rosa MGP. A simpler primate brain: the visual system of the marmoset monkey. Front Neural Circuits 8: 96, 2014.

Solomon SS, Chen SC, Morley JW, Solomon SG. Local and Global Correlations between Neurons in the Middle Temporal Area of Primate Visual Cortex. Cereb Cortex 25: 3182-3196, 2015.

Solomon SS, Tailby C, Gharaei S, Camp AJA, Bourne JA, Solomon SG. Visual motion integration by neurons in the middle temporal area of a New World monkey, the marmoset. J Physiol 23: 5741-5758, 2011.

Treue S, Martinez-Trujillo JC. Feature-based attention influences motion processing gain in macaque visual cortex. Nature 399: 575-579, 1999.

Uka T, DeAngelis GC. Contribution of middle temporal area to coarse depth discrimination: comparison of neuronal and psychophysical sensitivity. J Neurosci 23: 3515-3530, 2003.

Yu H-H, Chaplin TA, Davies AJ, Verma R, Rosa MGP. A Specialized Area in Limbic Cortex for Fast Analysis of Peripheral Vision. Curr Biol 22: 1-7, 2012.

Yu H-H, Chaplin TA, Egan GW, Reser DH, Worthy KH, Rosa MGP. Visually Evoked Responses in Extrastriate Area MT after Lesions of Striate Cortex in Early Life. J Neurosci 33: 12479-12489, 2013.

Yu H-H, Rosa MGP. A simple method for creating wide-field visual stimulus for electrophysiologyB: Mapping and analyzing receptive fields using a hemispheric display. J Vis 10: 1-16, 2010.

Zavitz E, Yu H-H, Row EG, Rosa MGP, Price Nicholas SC. Rapid Adaptation Induces Persistent Biases in Population Codes. J Neurosci 36: 4579-4590, 2016.

